# Genuage: visualizing and analyzing multidimensional point cloud data in virtual reality

**DOI:** 10.1101/2020.03.26.000448

**Authors:** Thomas Blanc, Mohamed El Beheiry, Jean-Baptiste Masson, Bassam Hajj

**Author notes:** contributed equally (Thomas Blanc, Mohamed El Beheiry). contributed equally (Jean-Baptiste Masson, Bassam Hajj).

## Abstract

The quantity of experimentally recorded point cloud data, such generated in single-molecule experiments, is increasing continuously in both size and dimension. Gaining an intuitive understanding of the data and facilitating multi-dimensional data analysis remains a challenge. It is especially challenging when static distribution properties are not predictive of dynamical properties. Here, we report a new open-source software platform, Genuage, that enables the easy perception, interaction and analysis of complex multidimensional point cloud datasets by leveraging virtual reality. We illustrate the benefit of the Genuage with examples of three-dimensional static and dynamic localization microscopy datasets, as well as some synthetic datasets. Genuage has a large breadth of usage modes, due to its compatibility with arbitrary multidimensional data types extending beyond the single-molecule research community.

## Main text

A consistent research trend today is to design and leverage experiments that generate massive databases. The complexity of data is equally increasing, raising a major concern on data presentation and interpretation, especially in early phase research. Several fields share the common feature of generating multidimensional point cloud data such as single-molecule localization microscopy [1], flow cytometry, transcriptomics and astrophysics to name a few. To tackle the interpretation of high-dimensional data, it has become common to seek low-dimensional embeddings of the N-dimensional data (N ≫ 3); typically, in to mostly two dimensions. In this format, it becomes possible to extract exploitable visual graphs [2], although at the price of lower fidelity to the ground truth and the concealment of interesting features.

A demonstrative example of this trend is found in single-molecule (SM) localization microscopy, where a single experiment can result in millions of individual 3D localizations [3]. New imaging approaches have furthermore extended localizations to encapsulate information complimentary to the position. These include temporal information (i.e. individual molecule trajectories) and molecular orientation [4], which notably grants new insights into how molecular arrangements influence global cellular architecture and function.

While automation and reproducibility of SM experiments remain a sought-after goal by many research groups, the fundamental challenge of extracting meaningful quantitative information from a single experiment persists. For example, extracting shapes from sparse SM point clouds, identifying a region of interest in a new experiment or asserting a domain boundary are complex tasks that resist standardization. Exploiting any geometric information stored in SM localizations is nontrivial; linking it to the evolution of dynamic properties (e.g. diffusion or drift) is particularly difficult as the statistical properties of point density may not be directly linked to their dynamical properties [5].

Recently, virtual reality (VR) has been re-introduced as a technology of interest for a vast number of applications due to the availability of low-cost consumer headsets and developer kits. The technology has already shown great promise for scientific research [6, 7]. Our conviction is that there is an immediate dual opportunity for VR in scientific research:

- Advanced representation and easy navigation of data to allow information to be disseminated rapidly
- Human-in-the-loop data treatment that mixes data interaction in VR to automated algorithms

With this motivation, we present Genuage, a VR-compatible open-source software platform compatible with any form of point cloud data, although we emphasize here its application to SM experiments. In complement to the approach described in ref [8] focusing on super-resolution, we extend the use of VR to dynamic data and higher dimensions. The design and architecture of Genuage were built around a simple basic concept that we see as essential in all scientific VR software: it is a 3D visualization vehicle. Specifically, VR serves to facilitate volumetric data interpretation by the user and can influence actions to take (i.e. human-in-the-loop data treatment) but it is an uncomfortable technology during extended uses. Genuage handles this issue with a dual visualization interface: a desktop mode for data loading and examination on a normal computer screen, and a VR mode for efficient data visualization, interaction and analysis. The desktop mode derives its general principles from the ViSP software by being ergonomic for users, allowing key parameters to be set while avoiding wearing the headset for long periods of time [9]. We emphasize that while progress in VR is rendering the technology more accessible, the experience remains far from comfortable and efficiency in processing the data is essential. The main characteristics of Genuage are summarized hereafter.

i. Genuage features an efficient means for loading and presenting in VR millions of data points within seconds (Supplementary Movie1). Point cloud data can be overlaid with other high-dimensional information such as molecular orientation, particle velocity, trajectories and color code in a robust and smooth way (Figure 1).
ii. The second main feature of Genuage, is a set of quantification tools specific to the VR mode. In VR, actions and measurements can be performed efficiently and precisely by the user in 3D (distances, angles, counting, local density calculator and histogram profiler) (Figure 1). In addition, the quantification set includes a selection tool for highlighting regions of interest for further analysis or for export (Figure 1, Supplementary movies 2-4). An automatically generated JSON metadata file records all the performed tasks for future sessions with Genuage and for post-treatment with other software. We provide a simple MATLAB script to illustrate how to read the JSON file and access the metadata.
iii. Genuage communicates with common software for data analysis such as Python and MATLAB, to exchange data sets after treatment (e.g. to be visualized in Genuage) or for further analysis (e.g. after selection and thresholding in Genuage). As such, applying already established quantification algorithms requires minimal effort in readapting the codes for specific programming language.
iv. In addition, Genuage offers a Bayesian analysis package derived from the InferenceMAP software [10] for live analysis of dynamic data in VR. In Figure 1 (Supplementary movie 5) we show an example of the trajectories of diffusing single molecules in the nucleus. The Suppelementary movie 5 shows the user interacting with 4D localization data with Genuage. The diffusion coefficient and forces are calculated on the selected trajectory or region of interest.

**Figure 1:**
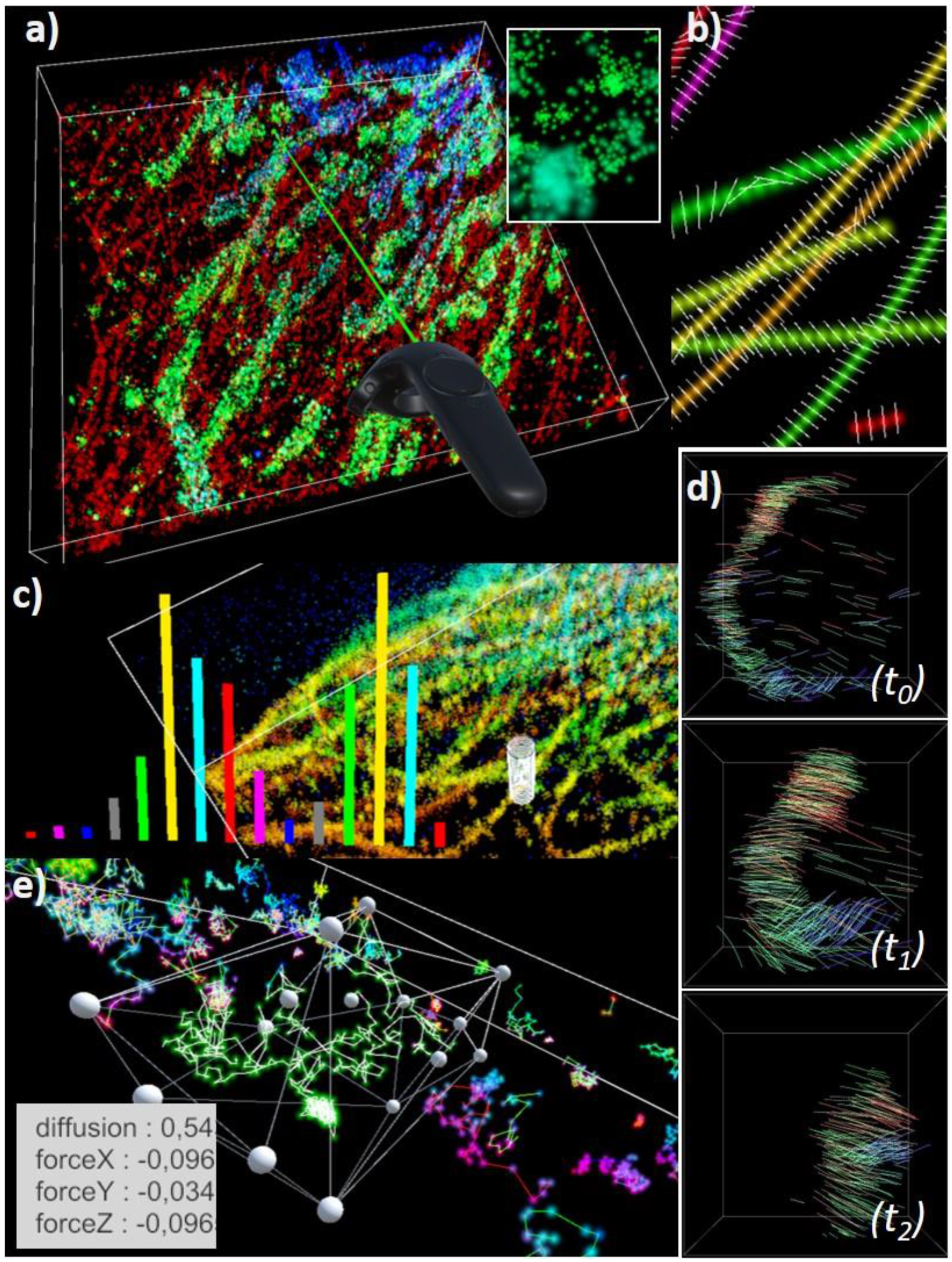
An overview of Genuage platform: a) visualization of 2 colors 3D super-resolution point cloud data in virtual reality, b) point cloud data can be overlaid with additional information such as molecular orientation, c) VR facilitates measurements and building histograms of molecular distributions in 3D, d) point cloud data with temporal information are explored, e) retrieving local dynamical properties by applying built-in algorithms on selected data points.

Genuage is an open-source software under a BSD license. Updated versions of the software will be regularly uploaded to <u>here</u>. We provide a detailed user manual describing the desktop and VR tools. While the examples presented in this paper are of single-molecule datasets, Genuage is adapted to other forms of multidimensional point cloud data generated synthetically or from other experiments. Supplementary videos show various examples of case studies and data manipulation in VR.

## Acknowledgments

We acknowledge funding from Fondation pour la recherche médicale – FRM– DEI20151234398 (B.H.), Agence National de la recherche ANR-19-CE42-0003-01, Labex Celtisphybio (ANR-10-LBX-0038), and Institut Curie (B.H.). We recognize the support by France-BioImaging infrastructure Grant ANR-10-INBS-04 Investments for the future (B.H.). We acknowledge the financial support of Agence pour la Recherche sur le Cancer (ARC Foundation) ARC (B.H.) and DIM ELICIT (B.H.). We acknowledge funding from the Pasteur Institute (J.B.M), the sponsorships of CRPCEN, Gilead Science and foundation EDF (J.B.M), the ANR-17-CE23-0016 TRamWAy (J.B.M), the INCEPTION project (PIA/ANR-16-CONV-0005, OG) (J.B.M), the programme d’investissement d’avenir supported by L’Agence Nationale de la Recherche ANR-19-P3IA-0001 Institut 3IA Prairie (J.B.M).

## Notes

https://github.com/Genuage/Genuage

