## Supplementary material for "Genuage: visualizing and analyzing multidimensional point cloud data in virtual reality"

### I. Genuage : Architecture and functionalities

Genuage is an open-source platform for visualization of n-dimensional point cloud data such generated in single-molecule super-resolution experiments. Genuage is coded in C# and built with the popular game engine Unity. Unity features (among numerous package) pre-built tools to render objects in a 3-dimensional space and deal with their movement. Genuage was tested with the following commercial VR headsets: with HTC VIve, Oculus Rift and Rift S. The compatibility is enhanced by the use of the VRTK library. Genuage is based on a modular object-oriented internal architecture which allows easy integration of new functionalities and modules in the code and also ensures the fluidity of the VR experience.

Genuage is based on two operating modes: a desktop mode and a virtual reality mode. This organization stems from the realization that although VR technologies have massively improved, spending a large amount of time in VR space is still not comfortable. VR experience should focus on the scientific perception and analysis of the data, while the desktop experience can be used for a first glimpse at the data, to setup parameters and preferences for the user.

**The desktop mode** provides means to setup the basic visual rendering of the point clouds. Users can load point cloud data and assign the input data columns to specific observable parameters. The observable parameters include 3D localizations, color code, time, trajectories for dynamical data and molecular 3D orientations. The values of the different columns can be thresholded and visualized directly. The color coding is performed with a predefined set of lookup tables. Users can change the size of the point clouds and the different axis scales. Moreover, multiple data files can be loaded and manipulated sequentially in the desktop interface. They can also be combined in the same visualization interface if needed. Exchanging data with Matlab and Python is possible in the desktop mode.

The **virtual reality mode** features new functionalities to visualized and analyse data. The VR mode in Genuage is a mean to an end. It serves the purpose of facilitating data interpretation and analysis. Several tools are provided. They are grouped into measurement tools (distances, angles, counting, and histograms), selection tools and visualization tools. Selections allow the user to isolate specific point clouds within any complicated mesh of 3D points followed by specific tasks such as exporting or analysis.

The different measurements and applied tasks can be saved into a JSON file to facilitate data re-opening. We provide a Matlab code to elucidate data accessibility by external software.

Besides, Genuage offers an ever-growing set of **analysis tools** such as density calculation in SR experiments and dynamic single-molecule data interpretation with Bayesian inference.

*Local density :* Genuage can calculate the density for all points in a cloud using a computational cost-effective approach. The space of the cloud is divided into several sub-regions and each point's density is calculated by counting all the points within the neighboring sub-regions that are within a radius set by the user. A new data column is generated with the point-wise calculated density values and could be assigned to a visual parameter such as color code for instance.

*Bayesian Inference*: Genuage implements and complements portions of the InferenceMAP [1] software to perform Bayesian Inference on selected trajectories of the data. Analysis is performed by sampling or optimizing the posterior distribution of parameters $P(U|T) \propto P_0(U).P(T|U)$, where U={D,**v**} represents the set of parameters, T the set of trajectories selected by the users, P(T|U) the likelihood of the trajectories knowing the parameters and $P_0$(U) the prior of the parameters. We used the likelihood introduced in [1,2] for diffusion and drift inference. Priors on the parameters can be flat (leading to a maximal likelihood inference), non-informative priors (see [3] for details) and conjugate priors (see [2] for examples). Depending on the source of the data, the user can have no positional noise, an homogenous noise in all spatial direction, a heterogenous noise with a planar noise and an out of plane noise. Genuage also features correction of diffusion due to motion blur (based on [4]).

Genuage being an open-source project: New tools will be continuously added, an extension of the manual and the supplementary featuring implementations and methods will be accessible on the GitHub: https://github.com/Genuage/Genuage.git.

The manual explains all the accessible features and analysis tools.

Several case studies are presented in the supplementary videos.

### II. Table of functionalities

A comparaison of the main tools and features emphasizing the complementarity between Genuage and two other tools for 3D single molecule data visualization and anlysis

| Functionalities | | VISP [5] | vLUME [6] | Genuage |
|---|---|---|---|---|
| **Interface** | Desktop mode | ✓ | × | ✓ |
| | VR mode | × | ✓ | ✓ |
| | Opening multiple files | ✓ into individual tabs | ✓ in the same working space | ✓ into individual tabs |
| | Accepted format | Text files | csv | .txt, .3d, any text files |
| **VR compatibility** | Headsets | - | HTC VIve, Oculus Rift and Rift S | HTC VIve, Oculus Rift and Rift S |
| **Visual adjustment** | Color adjustment | ✓ | ✓ | ✓ |
| | Axis scaling | × | ✓ | ✓ |
| | Point cloud size | ✓ | ✓ | ✓ |
| | Point cloud size by localization precision | ✓ | × | × |
| | Multi-color channels | ✓ | ✓ | ✓ |
| **Data input and visualization parameters** | Modular column assignement | × | × | ✓ |
| | Color code assigned to specific column | × | ✓ | ✓ |
| | 3D positions | ✓ | ✓ | ✓ |
| | Trajectories | × | × | ✓ |
| | Showing in time | × | ✓ | ✓ |
| | 2D Orientation | × | × | ✓ |
| | 3D Orientation | × | × | ✓ |
| | Thresholding | ✓ | ✓ | ✓ |
| **VR tools** | Clipping plane | - | ✓ | ✓ |
| | Selections | - | ✓ | ✓ |
| | Counting | - | × | ✓ |
| | Measuring distances | - | × | ✓ |
| | Histogram profiler | ✓in desktop mode | × | ✓ |
| | Annotation in VR | - | ✓ | × |
| | Data manipulation in VR | - | ✓ | × |
| **Built in analysis tools** | Point cloud local density calculation | ✓ | ✓ | ✓ |
| | Ripley's K function | × | ✓ | × |
| | Localization density | × | ✓ | × |
| | Largest and shortest distance | × | ✓ | × |
| | Diffusion coefficient | × | × | ✓$ |
| | Drift | × | × | ✓$ |
| **Reading/writing** | Exporting videos and figures | ✓imbeded | ✓imbeded | ✓Through external softwares |
| | Saving progress | × | ✓ | ✓Saving JSON file |
| | Exporting data | ✓ | ✓ | ✓exporting JSON file, exporting selections |
| | Communication with external software | × | × | ✓ |

$ Genuage provides up to 9 different methods for calculating the diffusion coefficient, and additional 10 methods for calculating  diffusion and drift.

### III. Data

Different 3D point cloud data sets are provided with the code for testing:

- Single-molecule localization data over 4 um depth generated by MultiFocus microscope. We provide localizations obtained by STORM super-resolution imaging of mitochondria (TOM-20 labled with Alexa 647) and another set for Tubulin structures (labled with Alexa 647).
- Single-particle localization data of injected beads in the nucleus of U2OS cells imaged in live with MFM. The corresponding trajectories are included.
- Simulated 3D point cloud trajectories with 3D orientation vectors

### IV. List of supplementary Video files

Supplementary videos can be downloaded here.

**Supplementary video S1:** General overview of Genuage interface. Example of loading and manipulating 2 colors point cloud data followed by distance measurement. The two colors are obtained by sequential super-resolution STORM imaging of fission-yeast cell-wall during division labeled with Alexa 647 followed by PALM imaging of tubuline fibers expressing tdEOS fluorescent protein in Mutlifocus microscopy (MFM) [7].

**Supplementary video S2:** Example of selection in VR of a complex 3D form of point clouds. The data is obtained by 3D STORM imaging of mitochondria using MFM.

**Supplementary video S3:** Clipping plane tool in VR to explore complex fiber network of point clouds generated by 3D super-resolution imaging of tubulin fibers of Hela cells in MFM.

**Supplementary video S4:** Using the profiler tool to measure the histogram of point clouds along a segment in 3D. The data correspond to 3D localizations obtained by STORM super-resolution imaging of TOM20 labeled with Alexa 647 dye. Such task is commonly performed to check the real resolution power of a super-resolution experience and to verify separation between point cloud features.

**Supplementary video S5:** Trajectory analysis performed on localizations of beads injected in the nucleus of U2OS cells and imaged by MFM. Local analysis and diffusion coefficient calculation in a dense 4D point cloud.

*OculusRift was used in videos 1and 5, HTC Vive was used in videos 2-4*
